# Graphic: Graph-Based Hierarchical Clustering for Single-Molecule Localization Microscopy

**DOI:** 10.1101/2020.12.22.423931

**Authors:** Mehrsa Pourya, Shayan Aziznejad, Michael Unser, Daniel Sage

**Affiliations:** Department of Electrical Engineering, Sharif University of Technology, Tehran, Iran; Biomedical Imaging Group, École polytechnique fédérale de Lausanne (EPFL), Switzerland

**Keywords:** Single molecule localization microscopy, spectral clustering, graph signal processing, multiscale methods, Delaunay triangulation

## Abstract

We propose a novel method for the clustering of point-cloud data that originate from single-molecule localization microscopy (SMLM). Our scheme has the ability to infer a hierarchical structure from the data. It takes a particular relevance when quantitatively analyzing the biological particles of interest at different scales. It assumes a prior neither on the shape of particles nor on the background noise. Our multiscale clustering pipeline is built upon graph theory. At each scale, we first construct a weighted graph that represents the SMLM data. Next, we find clusters using spectral clustering. We then use the output of this clustering algorithm to build the graph in the next scale; in this way, we ensure consistency over different scales. We illustrate our method with examples that highlight some of its important properties.

## 1. INTRODUCTION

Single-molecule localization microscopy (SMLM) is a very popular technique in super-resolution microscopy, with wide applications in cell biology [1]. This modality provides images that have a resolution in the order of 10-20 nm and is well suited to the study of structures at molecular scales inside a cell [2]. The data in SMLM are acquired by capturing thousands of images (a.k.a. frames), each one consisting of a small number of bright spots. After localizing each spot at every frame using computational techniques such as Gaussian fitting (see [3, 4] for detailed comparisons), one can render a compound high-resolution image on a discrete grid. It is noteworthy to mention that the raw output of the localization software is not a pixelized image, but rather a point-cloud (*i*.*e*., a list of spatial coordinates in nm). In addition, one often has access to extra information, such as some measure of uncertainty at each localized point, which is based on the number of associated emitted photons [5].

The analysis procedure often starts with a clustering of the acquired SMLM point-cloud. This is currently an active area of research [6], with density-based methods like DB-Scan [7, 8], persistence-based methods [9], tessellation-based methods [10, 11], and also Bayesian methods [12, 13, 14]. One common drawback of the existing methods is that they rely on some priors either on the shape of the structures of interest (*e*.*g*., particles of same size) and/or on the noise (*e*.*g*., uniform noise).

In this paper, we propose a novel graph-based method for hierarchical clustering (GrapHiC). It is a prior-free method that uses a multiscale grouping strategy. In our pipeline, we first build a graph based on a Delaunay triangulation of the data points. We then assign weights to the edges of the graph by modeling each node as a collection of bivariate Gaussian random variables. This allows us to incorporate information on the uncertainty of each localized spot. To perform clustering over the graph, we use a method called spectral clustering [15], which is well established in the field of graph signal processing [16]. After finding the first-level clusters, we create a new graph that summarizes the information of the previous step. By repeating this process, we infer a hierarchical structure on the input point-cloud. This multiscale strategy is motivated by the fact that SMLM data contain structural information that spans a wide range of scales: from cellular sub-compartments, to aggregates, all the way down to proteins. Our numerical experiments demonstrate the robustness and versatility of our scheme as well as its interesting hierarchical output.

### 1.1. Problem Formulation

The generic output of a 2D localization software is a point-cloud 𝒫= {**p**_1_, …, **p**_*M*_}, where **p**_*m*_ = (*x*_*m*_, *y*_*m*_), *m* = 1, …, *M* is the position of a localized point (in nm) and *M* is the number of points. The localization software often provides additional information; in this paper, we are particularly interested in the uncertainty *σ*_*m*_ and the number *N*_*m*_ of recorded photons associated to each point. The underlying hypothesis is that the *N*_*m*_ associated photons are i.i.d. realizations of a bivariate normal random variable 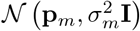. When the per-point photon count *N*_*m*_ is the sole output of the localization software, we translate it into an uncertainty through

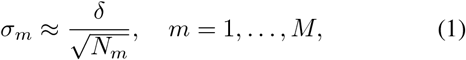

where *δ* is the size of the point-spread function of the microscope (typically, 250 nm).

Given the triplets (**p**_*m*_, *σ*_*m*_, *N*_*m*_) for *m* = 1, …, *M* as input data, our goal is to find a collection of injective labeling functions of the form

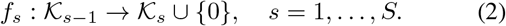

In (2), *S* denotes the number of scales and 𝒦_*s*_ is the set of aggregated points at the *s*th scale with the convention that 𝒦_0_ = 𝒫 is the input point-cloud. The label zero is specifically reserved for the data that are being discarded. We make it correspond to noisy localized points in the first scale and to comparatively small and isolated particles at higher scales.

## 2. METHOD

At each scale, our clustering method follows a three-step process: We first construct a graph from the input data; then, we preprocess the data by removing the isolated nodes; finally, we cluster the cleaned data by analyzing the eigenvalues of the corresponding Laplacian matrix (spectral clustering) [15]. By representing the clusters as aggregated points and repeating this procedure *S* times, we obtain a hierarchy of clusters. To facilitate the understanding of our method, we provide a running illustrative example in Figure 2.

**Fig. 1:**
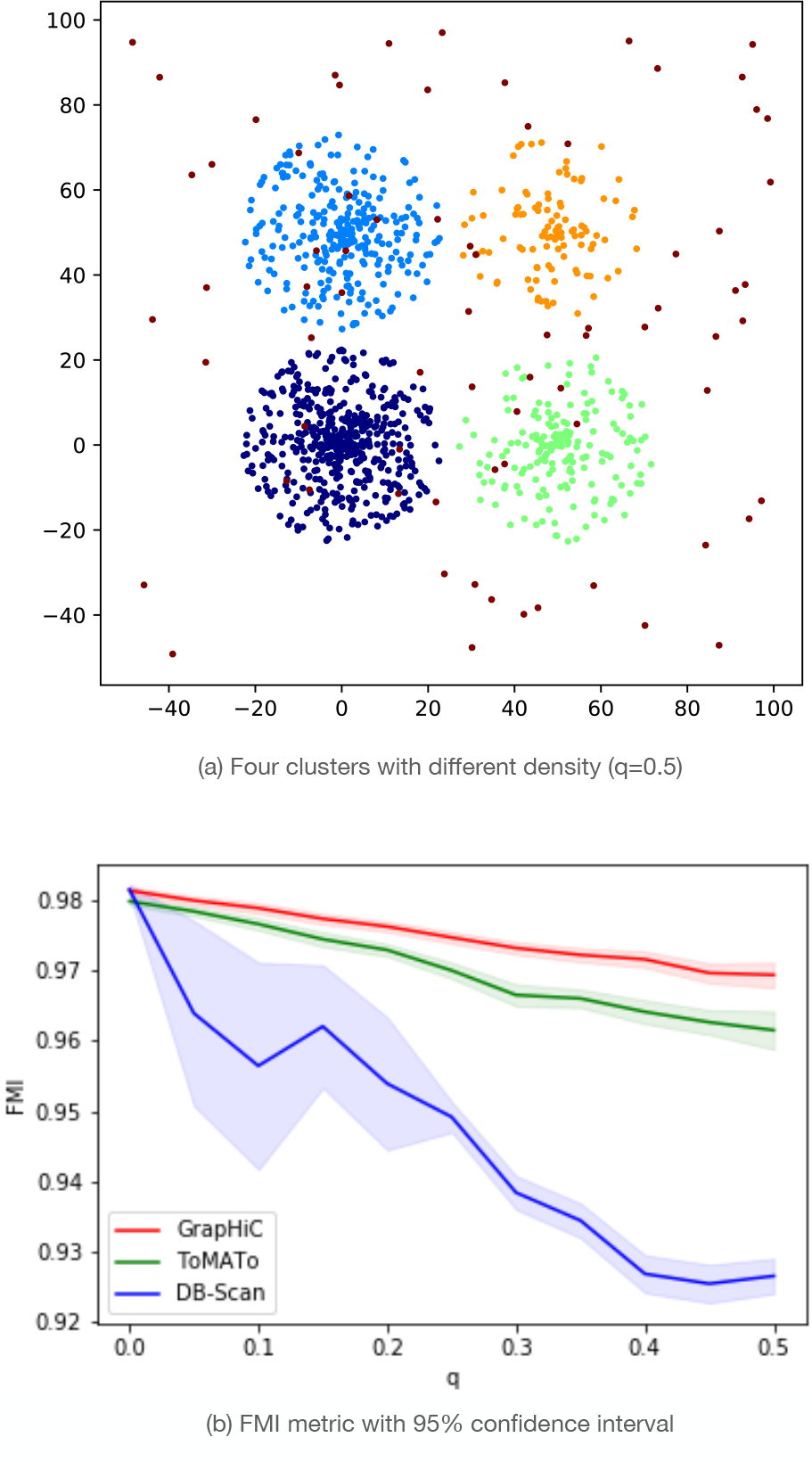
Performance of GrapHiC, ToMATo, and DB-Scan on the synthetic example. The results are aggregated over 50 runs.

**Fig. 2:**
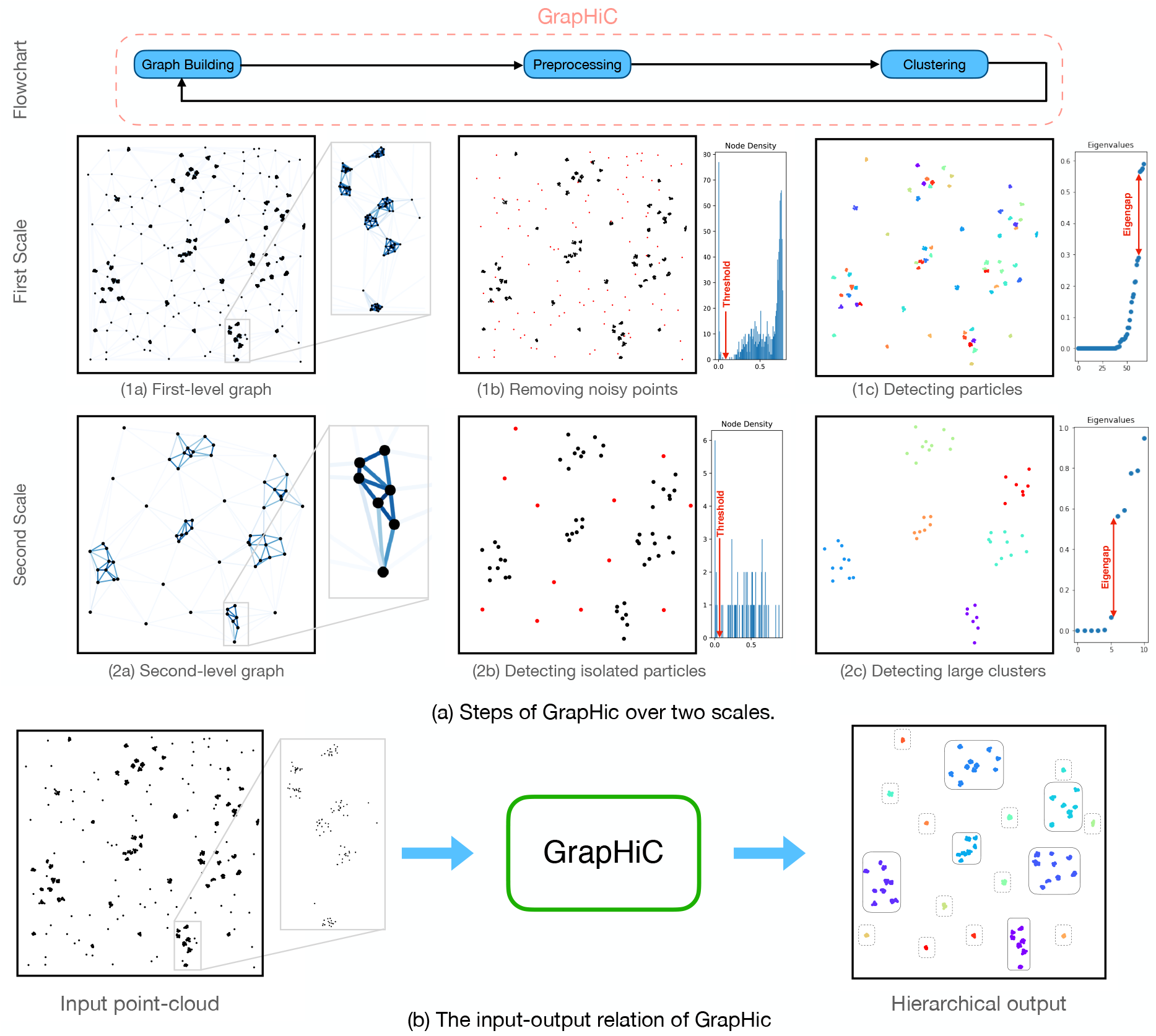
An illustrative example. We consider six large clusters, each with a random radius between 20 and 24. Three of them contain 10 and the other three contain 7 small particles. Besides, there are 12 dispersed particles as well (total number of 3 × 10 + 3 × 7 + 12 = 63 particles). Each particle has a random radius in [1.5, 3] and consists of 15 points. We also add 100 uniformly sampled points as the background noise.

### 2.1. Graph Construction

An undirected weighted graph 𝒢 = (𝒱, **W**) is a collection of nodes 𝒱 together with a square matrix **W** = [*w*_*m,n*_] ∈ ^ℝ | *𝒱* |*×*| *𝒱* |^ that is called the adjacency matrix of. In effect, *w*_*m,n*_ specifies the weight of the edge between vertices *v*_*m*_, *v*_*n*_ ∈ *𝒱*; it is set to zero if the nodes are disconnected.

#### 2.1.1. Nodes

At the *s*th scale, the input data are a collection of aggregated points, to each one we assign a bivariate normal distribution together with the number of recorded photons. Hence, the associated pairs of the form *v*_*m*_ = (𝒩 (**p**_**m**_, **Σ**_*m*_), *N*_*m*_), *m* = 1, …, | *𝒦*_*s*−1_| are the nodes of the graph at this scale.

#### 2.1.2. Edges

We perform a Delaunay triangulation on the position graph with the mean vector **p**_*m*_ assigned to the *m*th node. This gives us a preliminary connectivity (unweighted) graph. We then assign weights to its edges by computing the Gaussian similarity function [15]

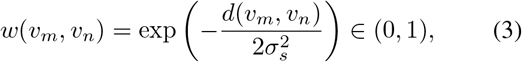

with then links *v*_*m*_ to *v*_*n*_. In (3), we set

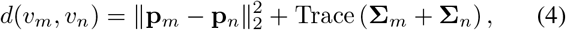

which is the expected value of the square Euclidean distance between two independent bivariate random vectors with distributions 𝒩 (**p**_*m*_, **Σ**_*m*_) and 𝒩 (**p**_*n*_, **Σ**_*n*_). The parameter *σ*_*s*_ controls the size of structures that we seek to find at the *s*th scale; indeed, the assigned weight for the edges that have an associated distance more than 3*σ*_*s*_ is approximately zero. Figure 2-(1a) provides an example of a weighted graph that we built in our pipeline.

### 2.2. Detection of Isolated Nodes

The first step at each scale is to remove from the graph the nodes that are isolated. To do so, we define the density *ρ*_*m*_ of the node *v*_*m*_ as

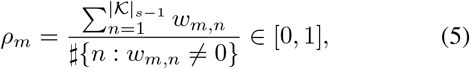

which is the average weight of the connected edges and measures how isolated a node is. The isolated nodes are then detected and removed from the original graph by applying a simple threshold to node densities, that is the node *v*_*m*_ will be removed if *ρ*_*m*_ is less than the threshold value. In Figure 2-(1b), we plot the density histogram and we highlight the desired threshold. There, we also show the cleaned point-clouds, where the isolated nodes are highlighted in red.

### 2.3. Spectral Clustering

Next, we apply the spectral-clustering method of [15] to the graph. Let us denote by 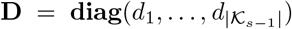, the degree matrix of the graph. It is a diagonal matrix whose *m*th entry *d*_*m*_ is equal to the sum of the edge weights that are connected to **v**_*m*_. The Laplacian matrix is then defined as **L** = (**D** − **W**) [16]. In spectral clustering, one analyzes the eigenvalues and eigenvectors of this matrix. It has been shown that the matrix **L** has exactly as many zero eigenvalues as the number of connected components of the corresponding graph. However, the existence of just a few weak links between the connected components gives rise to a set of nonzero (but small) eigenvalues. Hence, we estimate the number *K* of clusters by finding the first “significant” eigengap [15] (*e.g*., larger than 25% of the maximum peak) in the graph (see the eigenvalue plot in Figure 2-(1c)), by examining the difference between consecutive eigenvalues of the Laplacian matrix. Using this estimate, we then apply the well-known K-means algorithm to a collection of vectors of size *K*, where the *m*th node is represented by the vector of *m*th entries of eigenvectors that are corresponding to the *K* smallest eigenvalues.

### 2.4. Multiscale Clustering

So far, we have detailed our graph construction, preprocessing step, and clustering method at the *s*th scale. We note that the input data at this scale are the collection of pairs of the form *v*_*m*_ = (𝒩 (**p**_**m**_, **Σ**_*m*_), *N*_*m*_) for *m* = 1, …, | *𝒦* _*s*−1_|. In this section, we describe how to translate the output of the *s*th scale into a proper input at scale (*s* + 1).

Consider a generic cluster 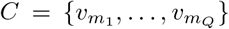 at the *s*th scale. We first estimate the mean vector, covariance matrix, and the photon count associated to *C* using the relations

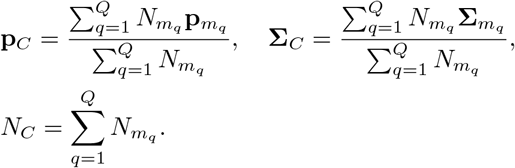

These estimates then allow us to represent *C* with the pair (𝒩 (**p**_*C*_, **Σ**_*C*_), *N*_*C*_). Doing so for clusters at the *s*th scale, we obtain the adequate pairs to be used in the next scale. As for the first scale, we simply assign the pair 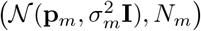 to the *m*th node of the graph for *m* = 1, …, *M*.

We demonstrate the results of a two-scale hierarchical clustering in Figure 2. The final display merges the outputs of the two scales to better infer the underlying hierarchical structure of the input data.

## 3. NUMERICAL RESULTS

In this section, we provide results of our experiments that are performed over both synthetic and real datasets.

### 3.1. Synthetic Example

We perform an objective comparison of our method with existing techniques over a dataset with a known ground-truth. The dataset consists of four round clusters with radius *r* = 23, each one having a different density. Precisely, the *k*th cluster is produced by uniformly sampling *n*_*k*_ = 500 × (1 − *q*)^*k*−1^ points for *k* = 1, 2, 3, 4, where *q* is a varying parameter. For a quantitative evaluation, we rely on the Fowlkes-Mallows index (FMI) [17] defined as.

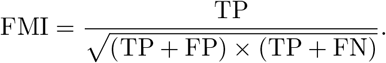

The result is depicted in Figure 1, where we clearly outperform DB-Scan [8] and ToMATo [9] for all values of *q*.

### 3.2. Real Dataset

For the second example, we consider now a region of the Paxilin dataset [14] and we apply our clustering pipeline for two scales. The results are depicted in Figure 3. As can be seen, our method successfully identified the elongated compartments that are present in this region.

**Fig. 3:**
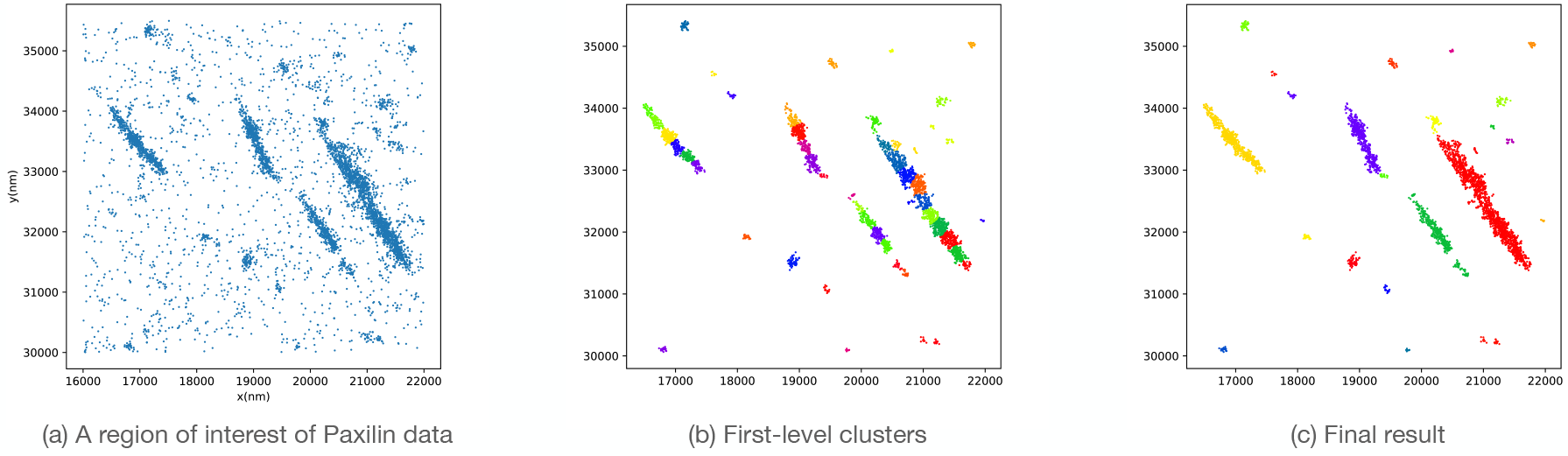
Output of GrapHiC on the Paxilin dataset. In the interest of clarity, we have removed small clusters with fewer than 5 points.

## 4. CONCLUSION

The proposed GrapHiC method is a graph-based hierarchical scheme for the clustering of SMLM data. In a multiscale strategy, we use spectral clustering on a graph that we build at each scale. Following several experiments, we demonstrated versatility and effectiveness of our method.

## 5. COMPLIANCE WITH ETHICAL STANDARDS

This article does not contain any studies involving human participants or animals performed by any of the authors.

### 6. ACKNOWLEDGEMENT

They also would like to thank Juliette Griffié and Thanh-An Pham for having fruitful discussions. They are also thankful to Ićiar Lloréns Jover for suggesting the acronym GrapHiC for our method.

There is no potential conflict of interest.

## REFERENCES

[1] L. Möckl and W. E. Moerner, “Super-resolution microscopy with single molecules in biology and beyond– essentials, current trends, and future challenges,” Journal of the American Chemical Society.

[2] M. Sauer, “Localization microscopy coming of age: from concepts to biological impact,” Journal of Cell Science, vol. 126, no. 16, pp. 3505–3513, 2013.

[3] D. Sage, H. Kirshner, T. Pengo, N. Stuurman, J. Min, S. Manley, and M. Unser, “Quantitative evaluation of software packages for single-molecule localization microscopy,” Nature Methods—Techniques for Life Scientists and Chemists, vol. 12, no. 8, pp. 717–724, 2015.

[4] D. Sage, T.-A. Pham, H. Babcock, T. Lukes, T. Pengo, J. Chao, R. Velmurugan, A. Herbert, A. Agrawal, S. Colabrese, A. Wheeler, A. Archetti, B. Rieger, R. Ober, G.M. Hagen, J.-B. Sibarita, J. Ries, R. Henriques, M. Unser, and S. Holden, “Super-resolution fight club: Assessment of 2D and 3D single-molecule localization microscopy software,” Nature Methods—Techniques for Life Scientists and Chemists, vol. 16, no. 5, pp. 387–395, 2019.

[5] H. Deschout, F. C. Zanacchi, M. Mlodzianoski, A. Diaspro, J. Bewersdorf, S. T. Hess, and K. Braeckmans, “Precisely and accurately localizing single emitters in fluorescence microscopy,” Nature Methods, vol. 11, no. 3, pp. 253–266, 2014.

[6] I. M. Khater, I. R. Nabi, and G. Hamarneh, “A review of super-resolution single-molecule localization microscopy cluster analysis and quantification methods,” Patterns, vol. 1, no. 3, pp. 100038, 2020.

[7] M. Ester, H. P. Kriegel, J. Sander, and X. Xu, “A density-based algorithm for discovering clusters in large spatial databases with noise.,” in Kdd, 1996, vol. 96, pp. 226–231.

[8] D. Bar-On, S. Wolter, S. Van De Linde, M. Heilemann, G. Nudelman, E. Nachliel, M. Gutman, M. Sauer, and U. Ashery, “Super-resolution imaging reveals the internal architecture of nano-sized syntaxin clusters,” Journal of Biological Chemistry, vol. 287, no. 32, pp. 27158–27167, 2012.

[9] J. A Pike, A. Khan, C. Pallini, S. G Thomas, M. Mund, J. Ries, N. S Poulter, and I. B Styles, “Topological data analysis quantifies biological nano-structure from single molecule localization microscopy,” Bioinformatics, vol. 36, no. 5, pp. 1614–1621, 2020.

[10] F. Levet, E. Hosy, A. Kechkar, C. Butler, A. Beghin, D. Choquet, and J. B. Sibarita, “Sr-tesseler: a method to segment and quantify localization-based super-resolution microscopy data,” Nature Methods, vol. 12, no. 11, pp. 1065–1071, 2015.

[11] L. Andronov, I. Orlov, Y. Lutz, J. L. Vonesch, and B. P. Klaholz, “Clustervisu, a method for clustering of protein complexes by Voronoi tessellation in super-resolution microscopy,” Scientific Reports, vol. 6, no. 1, pp. 1–9, 2016.

[12] J. Griffié, M. Shannon, C. L. Bromley, L. Boelen, G. L. Burn, D. J. Williamson, N. A. Heard, A. P. Cope, D. M. Owen, and P. Rubin-Delanchy, “A Bayesian cluster analysis method for single-molecule localization microscopy data,” Nature Protocols, vol. 11, no. 12, pp. 2499, 2016.

[13] J. Griffié, L. Shlomovich, D. J. Williamson, M. Shannon, J. Aaron, S. Khuon, G. L. Burn, L. Boelen, R. Peters, A. P. Cope, E. Cohen, P. Rubin-Delanchy, and Owen D. M., “3D Bayesian cluster analysis of super-resolution data reveals LAT recruitment to the T cell synapse,” Scientific Reports, vol. 7, no. 1, pp. 1–9, 2017.

[14] H. Deschout, I. Platzman, D. Sage, L. Feletti, J. P. Spatz, and A. Radenovic, “Investigating focal adhesion substructures by localization microscopy,” Biophysical Journal, vol. 113, no. 11, pp. 2508–2518, 2017.

[15] U. von Luxburg, “A tutorial on spectral clustering,” Statistics and Computing, vol. 17, no. 4, pp. 395–416, 2007.

[16] A. Ortega, P. Frossard, J. Kovačević, J. Moura, and P. Vandergheynst, “Graph signal processing: Overview, challenges, and applications,” Proceedings of the IEEE, vol. 106, no. 5, pp. 808–828, 2018.

[17] E. B. Fowlkes and C. L. Mallows, “A method for comparing two hierarchical clusterings,” Journal of the American Statistical Association, vol. 78, no. 383, pp. 553–569, 1983.

